# Pneumococcal Extracellular Vesicles Mediate Horizontal Gene Transfer via the Transformation Machinery

**DOI:** 10.1101/2023.12.15.571797

**Authors:** Sarah Werner Lass, Shaw Camphire, Bailey E. Smith, Rory A. Eutsey, Jojo A. Prentice, Saigopalakrishna S. Yerneni, Ashni Arun, Andrew A. Bridges, Jason W. Rosch, James F. Conway, Phil Campbell, N. Luisa Hiller

## Abstract

Bacterial cells secrete extracellular vesicles (EVs), the function of which is a matter of intense investigation. Here, we show that the EVs secreted by the human pathogen *Streptococcus pneumoniae* (pneumococcus) are associated with bacterial DNA on their surface and can deliver this DNA to the transformation machinery of competent cells. These findings suggest that EVs contribute to gene transfer in Gram-positive bacteria, and in doing so, may promote the spread of drug resistance genes in the population.

**Significance:** This work extends our understanding of horizontal gene transfer and the roles of extracellular vesicles in pneumococcus. This bacterium serves as the model for transformation, a process by which bacteria can take up naked DNA from the environment. Here we show that extracellular vesicles secreted by the pneumococcus have DNA on their surface, and that this DNA can be imported by the transformation machinery facilitating gene transfer. Understanding EV-mediated gene transfer may provide new avenues to manage the spread of antibiotic drug resistance.

## Introduction

There is a high degree of genomic diversity and plasticity across strains of *Streptococcus pneumoniae* (pneumococcus or Spn), where the pangenome extends well beyond the genes found in any single strain (1, 2). This plasticity is driven by widespread horizontal gene transfer (HGT) between pneumococcal strains and strains of related species (3). HGT in pneumococcus can occur via transduction, conjugation, and transformation. Accordingly, its pangenome encodes a high number of phages, integrative conjugative elements, and evidence of transformation events (1, 4–7).

Pneumococcus is the paradigm of natural competence. Evidence of gene transfer and participating molecules dates back to Griffith’s “transforming principle” in 1928 (8) and the first bacterial quorum sensing factor described in 1965 (9). Transformation is traditionally defined as the uptake of naked DNA from the environment, often derived from lysed pneumococcal cells (10). Transformation is initiated by the autoinducing Competence Stimulating Peptide (CSP) (11). CSP activates a two component system (ComD and ComE), leading to changes in the expression of over 5% of genes in the pneumococcal genome (12, 13). A critical phenotypic consequence of CSP induction is the assembly of the transformasome, a multi-protein complex that imports single-stranded DNA from the immediate extracellular environment and delivers it to RecA for recombination (14, 15). In this manner, CSP initiates the process by which cells can take up such DNA, a state known as competence. Transformation is active during chronic mucosal infection and colonization events (7, 16).

The hypothesis that extracellular vesicles (EVs), also known as membrane vesicles or outer-membrane vesicles in Gram-negative species, may serve as vehicles for HGT was put forward decades ago. In 1989, Judd and colleagues observed DNA within EVs from *Neisseria gonorrhoeae* (referred to as membrane blebs) and demonstrated the transfer of plasmid DNA from the lumen of EVs to cells of the same species (17). Subsequent investigations involving other Gram-negative species, such as *Escherichia coli*, *Pseudomonas aeruginosa*, *Acinetobacter baylyi*, and *Acinetobacter baumannii*, revealed that EVs from these bacteria can carry either plasmid or chromosomal DNA (18–21). Intra-species DNA delivery via EVs was observed for *E. coli* and *Acinetobacter* (18, 21). Further, EV-mediated inter-species DNA delivery was shown from *E. coli* to *Salmonella enterica* serovar Enteritidis and from *A. baylyi* to *E. coli* (18). In *A. baylyi*, intra-species transfer of DNA required an intact competence system, suggesting that EV-mediated HGT may not be a passive process for the recipient cell (21). In contrast to Gram-negative EVs, much less is known about EVs produced by Gram-positive bacteria. EVs secreted by viable Gram-positive bacteria must traverse the thick cell wall, a process that is not well understood (22).

Multiple studies on the pneumococcus have demonstrated that this bacterium secretes EVs (23–27). Pneumococcal EVs (pEVs) have been shown to be internalized by host cells, and to interact and influence multiple components of the mammalian immune response including the complement system, dendritic cells, macrophages, and neutrophil extracellular traps (24–26, 28). However, it remains unclear whether pEVs interact with and alter the physiology of the source population.

In this study, we establish that pEVs serve as DNA donors to pneumococcal cells. Our study demonstrates that genomic DNA associates with the external surfaces of pEVs. This DNA can serve as a source for transformation but requires CSP signaling and transformation machinery on the recipient cell. Thus, we propose that pEVs transport DNA and deliver it to the transformation machinery, contributing to pneumococcal gene transfer. This model suggests that the biogenesis, secretion, transport, or uptake of pEVs may influence the dynamics of transformation during infections.

## Results

### Purification of pneumococcal EVs (pEVs)

We purified pEVs from cultures at late-log phase by size exclusion chromatography (SEC) (28). This yielded 1 mL of purified pEVs from 500 mL of culture, containing approximately 4.8×10^10^ vesicles/mL and a corresponding protein content of approximately 21 µg/mL. Nanoparticle tracking analysis (NTA) of pEVs suggested they span a wide range of sizes, from 25-400 nm in diameter, with a median size of 129 nm (**SFig.1A**).

To visualize the pEVs, we subjected the SEC-purified fraction to cryo-electron microscopy (cryo-EM). The images revealed round vesicles with a clear membrane bilayer (**SFig.1B, C**). Vesicles displayed heterogeneity in size, consistent with the range measured by NTA. The external surfaces of the pEVs also appeared heterogeneous. Some pEVs displayed a smooth surface, while others appeared textured, as if decorated with macromolecules (often referred to as an EV biocorona (29)). Several images captured pEVs that appear to be undergoing fusion or fission, as well as EVs encapsulating other pEVs (doublets or triplets) (**Fig. S1**). Together, the NTA and cryo-images confirm the presence of pEVs in our SEC fraction and reveal their morphological heterogeneity.

**SFig.1:**
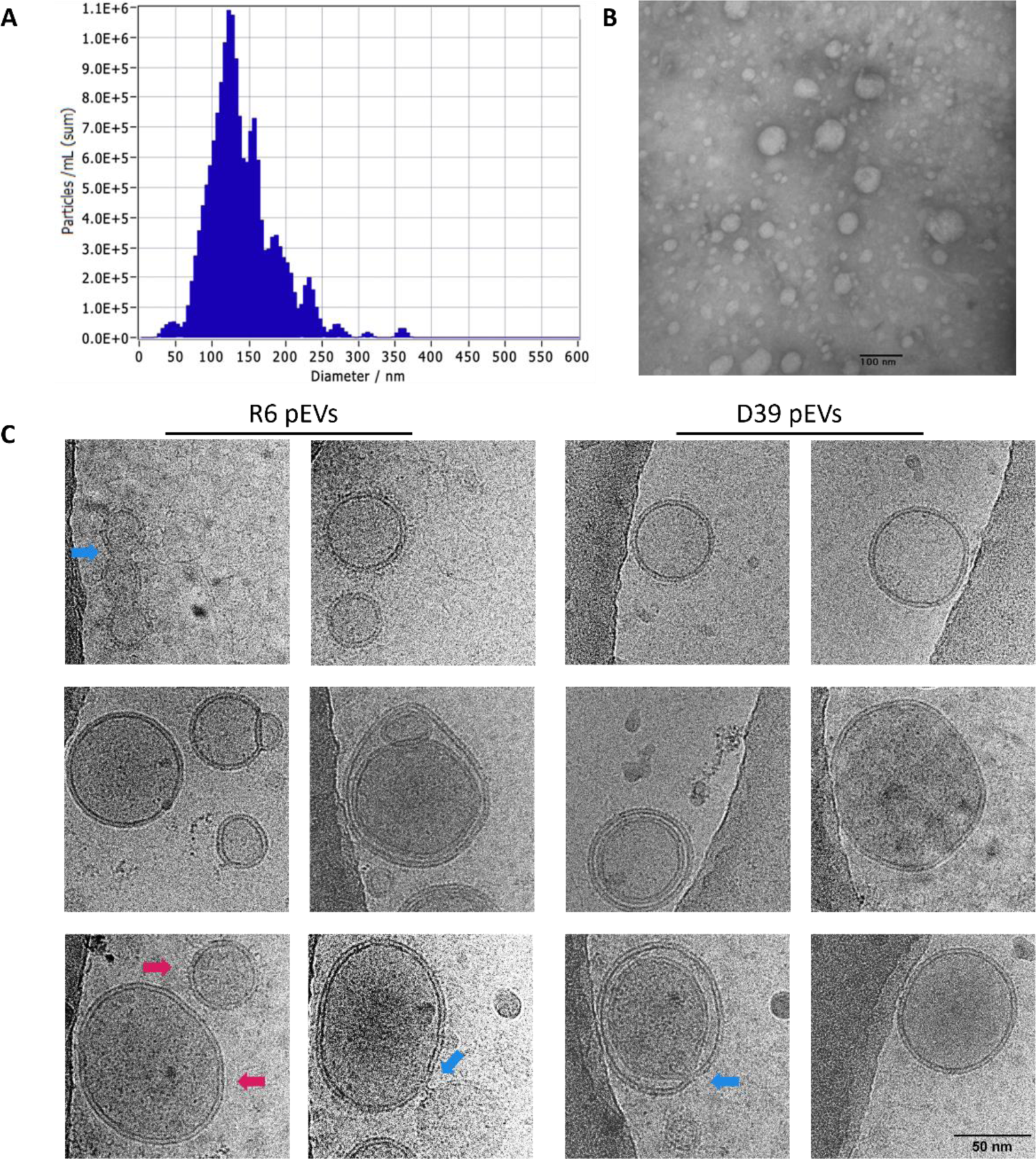
(**A**) Representative nanoparticle tracking analysis of pneumococcal extracellular vesicles isolated from R6 (produced by the software ZetaView). (**B**) Representative transmission electron micrograph of pEVs from R6, scale bar 100 nm. (**C**) Images selected from cryo-electron micrographs of pEVs from R6 and D39. The first row displays representative features observed across most images. Magenta arrows indicate two pEVs representing either smooth or textured surfaces. Blue arrows indicate apparent fusion or fission events (rare in our set). Doublet and triplet pEVs are also relatively rare in our set. All images are the same scale (scale bar of 50nm in final image).

### DNA associates with the extracellular surface of Pneumococcal EVs

Since EVs from multiple organisms carry DNA as cargo both in the lumen and on external surfaces, we first investigated whether DNA is associated with pEVs (21, 30–32). To answer whether DNA was present in our pEV samples, we used pEVs as a template for a polymerase chain reaction (PCR) and successfully amplified a pneumococcal-specific DNA sequence (**Fig.1A**). This suggested that DNA is associated with pEVs.

Next, we investigated the localization of DNA in relation to the pEVs. We reasoned that DNA could be in the luminal compartment of pEVs or could be associated with the external surface, and that DNA in either location would be an available template for PCR as thermal cycling ruptures the pEVs. To differentiate between external and luminal DNA presence, we sequentially treated pEVs with a non-membrane permeable DNase (Turbo DNase), inactivated that DNase, then treated with a detergent to lyse pEVs and expose luminal DNA. Untreated pEVs are associated with DNA, as measured by PCR amplification. However, once samples are exposed to a DNase that targets their surface, we can no longer amplify DNA by PCR, suggesting that detectable DNA is on the pEV surface (**Fig. 1A, lanes 1 versus 2**). This loss of signal is not a result of residual DNase activity in the sample or of loss of polymerase activity, as spiking the sample with DNA rescued the PCR signal (**Fig. 1A, lanes 3-4**). To ensure that intra-EV DNA was exposed, we lysed the pEVs after DNase digestion and inactivation to increase accessibility of any luminal DNA as a template for PCR. This sample has no PCR signal, suggesting there is no detectable DNA in the pEV lumen. This lack of signal is not a result of Triton interference with PCR amplification, as evidenced by spiking the sample with DNA (**Fig. 1A, lanes 5-6**). While we could not identify luminal pEV DNA, there remains the possibility of low luminal pEV DNA concentrations identifiable using more sensitive experimental approaches. Taken together, we conclude that DNA is associated with the outer surface of pEVs and not present within the pEV lumen.

To further evaluate the association between pEVs and DNA we performed cryo-EM and simultaneous two camera imaging using a spinning disc confocal microscope on pEVs. The cryo-EM was performed on pEVs isolated from unencapsulated strain R6 and its encapsuled ancestor D39. The cryo-EM images revealed nucleic acid strands surrounding the pEVs in both strains (**Fig. 1B**). Using simultaneous two camera imaging, we measured the extent of co-localization and co-diffusion of DNA and lipids in the pEV sample for strain D39. For visualization, we used the non-membrane permeable dsDNA stain PicoGreen, and for the pEV membrane we used DiD lipophilic dye. Our analyses captured the co-localization and the synchronized movement of both molecules in the pEV samples, demonstrating that DNA is associated with pEVs (**Fig. 1C-E**). We conclude that DNA is associated to the outer surface of pEVs generated from both encapsulated and non-encapsulated cells.

To characterize the length and source of the DNA on the pEVs, we used PCR and whole genome sequencing. Using primers that target multiple locations on the pneumococcal genome, we amplified DNA of 1, 3, 5 and 7 Kb, suggesting that individual fragments of pEV DNA can be at least of this size (**Fig 1A, SFig.3A-B**). Next, we investigated whether DNA on pEVs corresponds to the whole genome or is enriched for specific sequences. To this end, we purified DNA from pEVs and their parental strains (**SFig.3C**) and submitted these samples to whole genome sequencing (WGS). We used pEVs from three *Spn* strains: R6, a lab adapted strain; B1599, a classic non-typeable strain commonly associated with eye infections (33), and ATCC700669-NC, a strain from the pandemic PMEN1 lineage with a naturally occurring deletion within the capsular locus. Our sequencing results revealed at least 98% coverage of the genome on the pEVs from each strain (**Fig.1B**). We conclude that pEV DNA is broadly representative of the parent bacterium.

**Fig.1:**
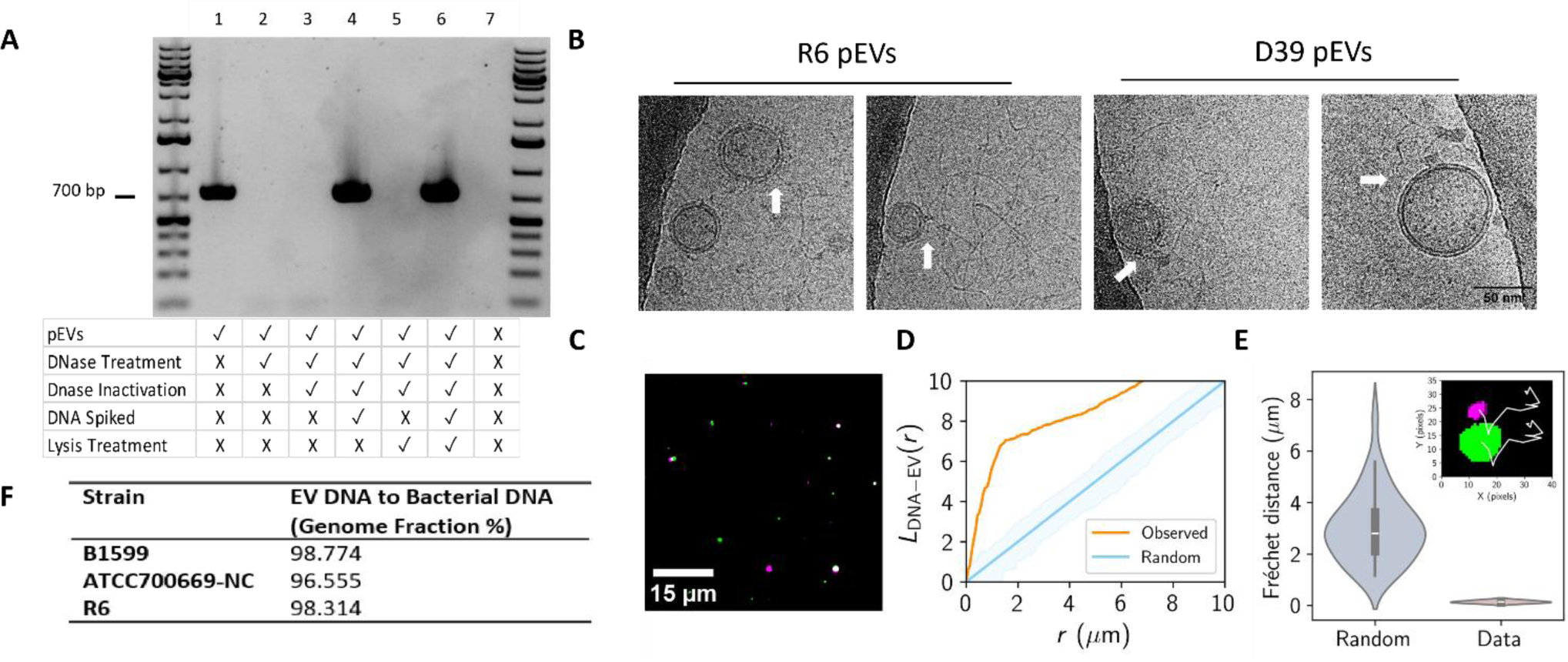
Genomic DNA is present on the surface of pEVs. **(A)** PCR analysis of pEV DNA localization. pEVs underwent a series of treatments to determine the location of DNA (representative gel, n=3). At each treatment step, including the untreated pEVs, a sample was saved for PCR amplification and subsequent gel electrophoresis. The rows below each lane indicate each sample’s treatment. DNase treatment was performed with 1 U Turbo DNase at 37 °C for 30 minutes. DNase inactivation was accomplished with 5 µM EDTA (f.c.) for 10 minutes at 75 °C. Samples that were spiked with DNA had 50 ng of genomic DNA added. Lysis treatment was performed with 1% triton (f.c.) at 65 °C for 10 minutes The negative control in the last lane includes only 1x PBS, the buffer used for PEV elution from SEC. Primers targeted the gene *gapdh*. First and last lanes include the GeneRuler 1 kb Plus DNA Ladder (Invitrogen). **(B)** Images selected from cryo-electron micrographs of pEVs from R6 and D39 that display nucleic acid strands surrounding pEVs. White arrows indicate point of association between pEVs and nucleic acid strands. All images are the same scale (scale bar in final image). **(C-D)** Spatial analysis of DNA and pEV co-localization. **(C)** Representative image of DNA particles, false colored in green, and pEV particles, false colored in magenta. pEV sample was treated with PicoGreen and DiD to label DNA and the pEV membrane, respectively. Scale bar is indicated. **(D)** Ripley’s cross-L function for clustering of simulated random (blue) and observed (orange) pEV particles to DNA particles as a function radius, r. Data represent n=4 independent fields of view of DNA and pEV molecules (n=175 pEV particles and n=477 DNA particles). Shading represents the acceptance region for a hypothesis test of complete spatial randomness, with significance level 5%, using the envelopes of the L-functions of 1000 simulations. **(E)** DNA and pEV co-diffusion. Fréchet distances computed for simulated (Random) and experimental (Data) of pEV-DNA particle trajectories. Data represent n=4 pEV-DNA pairs. Random Fréchet distance determined using diffusion coefficients for each pair determined in SFig 2. Inset: representative trajectories (white) from a DNA (green) and pEV (magenta) pair. **(F)** Sequencing of DNA isolated from pEVs and their parent bacterium from three unencapsulated pneumococcal strains (n=1 per strain).

**SFig.2:**
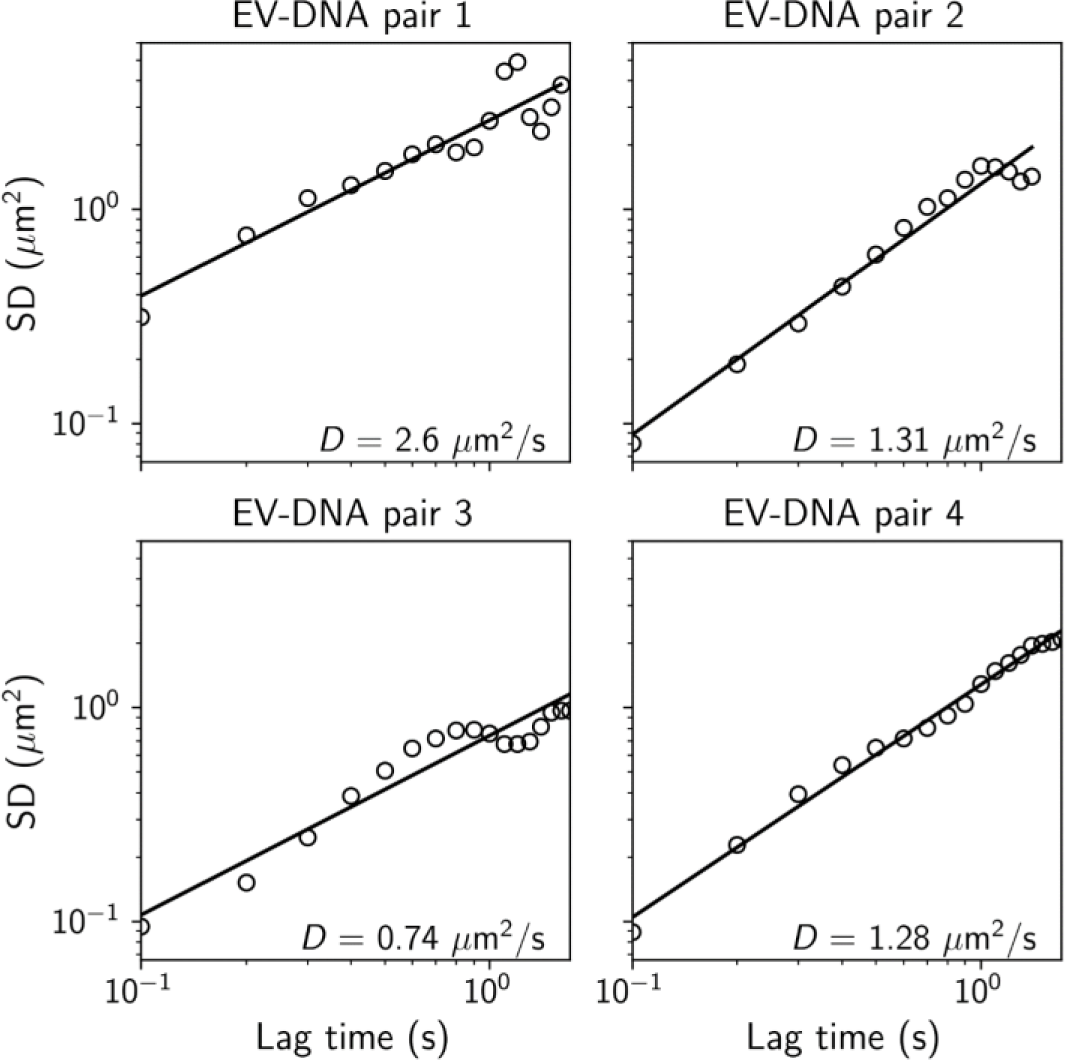
pEV particles follow Brownian trajectories. Points represent computed squared displacements (SD) as a function of lag time for tracked pEV particles. Lines represent linear fits to the data in log-space for each pEV particle. D = Diffusion coefficients. **SVideo.1**: Co-localization and co-diffusion of pEV and DNA particles. DNA particles are false colored in green (left panel), pEV particles are false colored in magenta (middle panel), and the merge is present in the right panel. pEV sample was treated with PicoGreen and DiD to label DNA and the pEV membrane, respectively. Scale bar is indicated.

To quantify DNA on the pEVs, we employed PicoGreen dye and qPCR. The concentrations of DNA present in pEV samples were highly variable. pEVs purified from strain R6 displayed a range of 0.1-3.2 ng/µL and an average 1.1 ng/µL of DNA. We also tested the parallel values for the encapsulated ancestor of R6, strain D39. D39 pEV samples displayed a range of 3.5-6.0 ng/µL and an average 4.8 ng/µL of DNA. Finally, we performed qPCR on pEVs for further quantification and confirmation of DNA presence. The measurements are in a similar range as PicoGreen, confirming that pEV samples carry between 0.4-7.7 ng/µL of DNA. All DNA measurements, when normalized to total pEV particle number, were not significantly different across measurement method (PicoGreen vs qPCR) or strain (R6 vs D39) (**SFig.3D**). We estimate that the DNA carried by pEVs represents an average of 1 genome equivalent per 16 pEVs, with a range spanning 1 genome equivalent per 4-324 pEVs depending on the purification.

**SFig.3:**
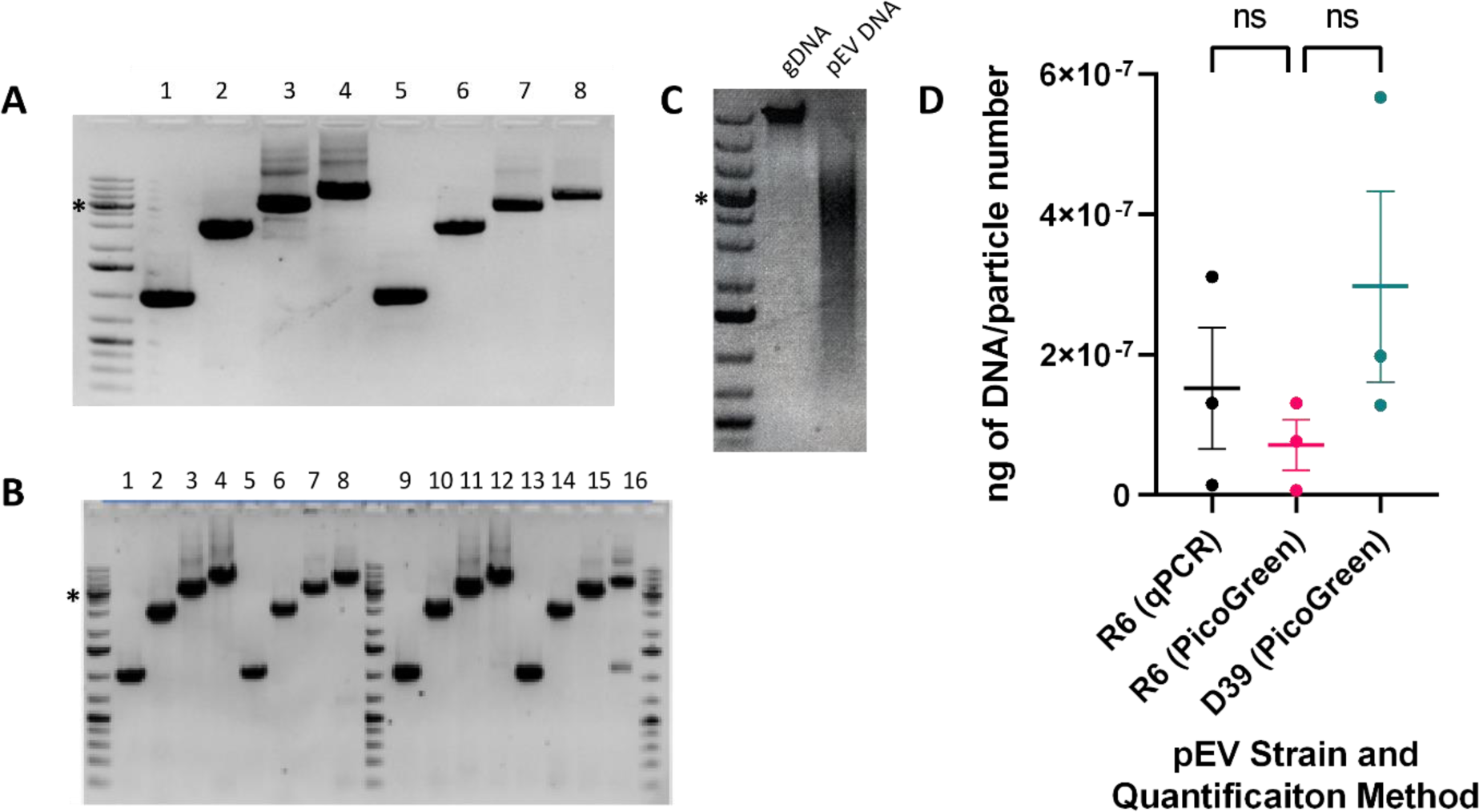
**(A-B)** PCR amplification of 1, 3, 5, and 7 kb DNA fragments from R6 pEVs and genomic DNA. **(A)** PCR primers targeted to gene spr1608. Left lane is the MW standard; lanes labeled 1-4 are positive control where genomic DNA was used as template, and lanes 5-8 used pEVs as template. PCR was performed on two independent sets of vesicles. **(B)** PCR primers targeted to gene spr0001 (left half of the gel) and spr0065 (right half of the gel). Lanes 1-4 and 9-12 are positive control where genomic DNA was used as template, lanes 5-8 and 13-18 used pEVs as template. **(C)** Gel electrophoresis profile of DNA purified from bacterial culture (gDNA) or SEC-purified pEVs (pEV DNA). Asterisk (*) indicates 5,000 base pair marker on the GeneRuler 1 kb Plus DNA Ladder (Invitrogen). **(D)** Quantification of pEV DNA by qPCR and PicoGreen. qPCR was performed on three R6 pEV samples and each point represents four technical replicates. PicoGreen staining was performed on the same three R6 pEV samples and three D39 pEV samples. These data are normalized to the number of pEVs in each sample as measured by NTA. One-way ANOVA compared to R6 (PicoGreen), ns=p>0.5.

### pEVs serve as a source of DNA for transformation

We next sought to ascertain whether DNA associated with pEVs facilitates gene transfer. To this end, we used pEVs from an R6 derivative that encodes spectinomycin-resistance (R6-SpecR) to be used as a marker for gene transfer. When pEVs alone were mixed with spectinomycin-sensitive R6 cells (R6-SpecS), we did not observe gene transfer, that is, there were no spectinomycin-resistance cells (**Fig. 2**). These data suggest that, under these conditions, pEVs were not internalized into the pneumococcal cells in a manner that delivered DNA capable of recombination.

We reasoned that DNA on the pEV surface may be accessible to the transformation machinery of competent pneumococcal cells. To test this hypothesis, we evaluated the ability of competent R6-SpecS to take up DNA from pEVs produced by R6-SpecR and used R6-SpecR genomic DNA as a positive control. Competence was induced by the addition of CSP. This experiment yielded spectinomycin resistant transformants (**Fig. 2**). To confirm the colonies were indeed transformants, three transformant colonies were selected from each plate, sub-cultured in rich media, grown to late-log phase, and an aliquot was used as a DNA template for PCR (**SFig. 4A**). Every transformant tested was positive for the gene encoding spectinomycin resistance.

**Figure 2.**
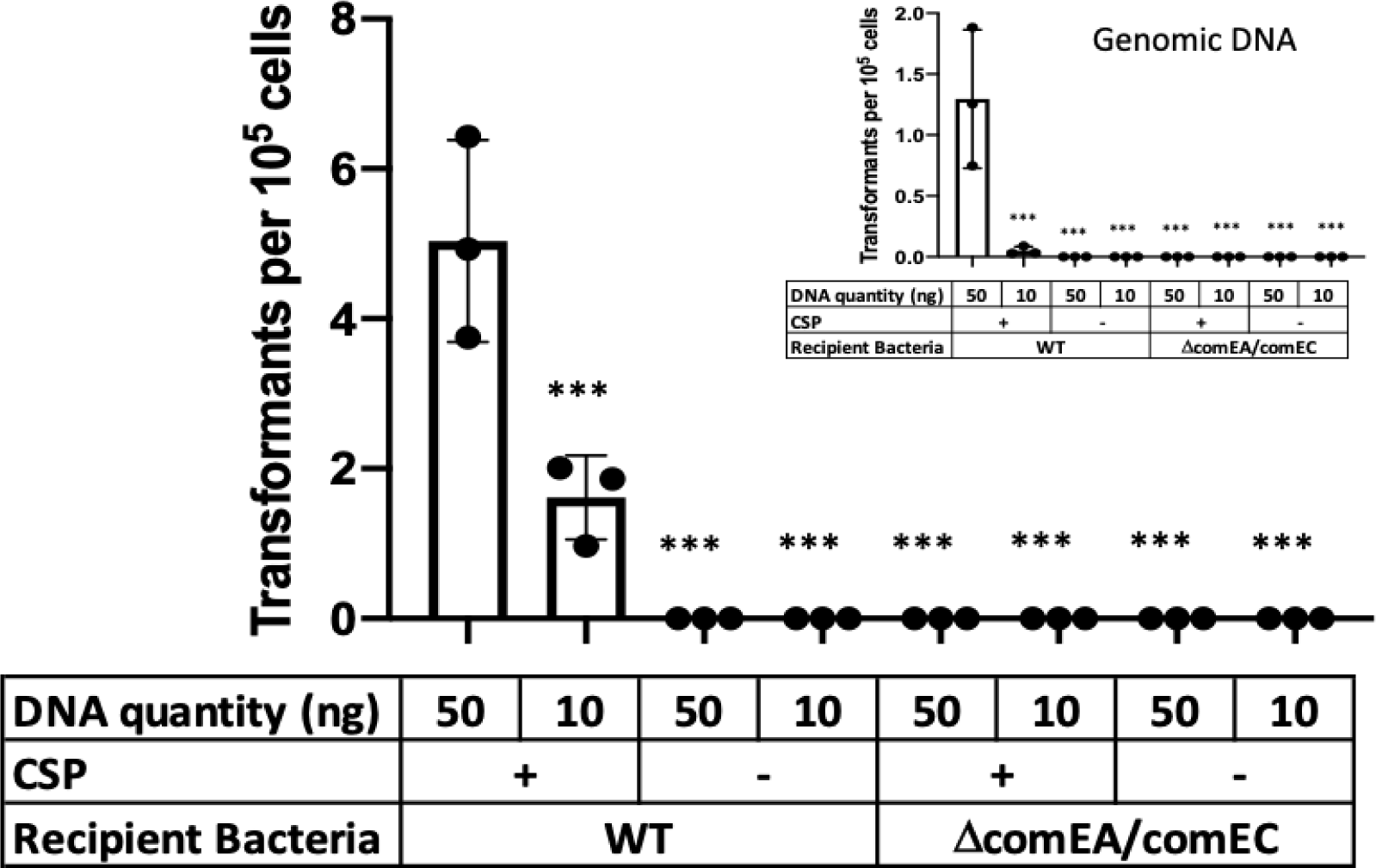
pEVs serve as a source of DNA for competent Spn cells. R6 cells (SpecS background) were exposed to pEV DNA (main figure) or genomic DNA (inset) from a R6-SpecR strain. Transformations were performed with and without CSP. Further, we tested recipient cells that do not encode a functional transformasome (Δ*comEA*/Δ*comEC*). Given the variability in efficacy between pEV batches, we did not draw conclusions about transformation efficiencies between pEVs or genomic DNA-mediated transformations. We propose that the range of transformation efficiencies reflects pEVs heterogeneity. Bars represent mean ± SEM with dots overlayed within a bar representing a data point from each independent experiment (n ≥ 3, *** adjusted p-value < 0.0001 for Dunnett’s multiple comparison test).

While some pneumococcal strains are non-encapsulated, most encode a capsule (34). To test whether the capsule represents an additional barrier for DNA transfer from pEVs to the transformasome, we tested whether pEVs generated from the encapsulated strain D39 could also serve as a DNA source for transformation. To this end, we isolated pEVs from D39-SpecR and exposed them to D39-SpecS cells, in the presence and absence of CSP and with genomic DNA from D39-SpecR as a positive control. We found that, akin to R6, pEVs as well as genomic DNA serve as a source for gene transfer and this process requires CSP (**SFig.4B**). We conclude that pEVs, from both non-encapsulated and encapsulated strains, can serve as a source of DNA for recombination in competent cells.

Competence influences many pneumococcal processes (13). To directly test the role of the transformation machinery in uptake of pEV DNA, we generated a strain with deletions in *comEA* and *comEC* in the R6-SpecS background (*ΔcomEA/comEC*), as these genes are required for a functional transformasome (35). We found that this strain is not transformed by pEVs, even in the presence of CSP (**Fig. 2**). We conclude that DNA from pEVs enters the cells via the transformasome of competent cells. Overall, we conclude that pEVs carry DNA on their surface, and this DNA can be transferred to competent cells in a manner dependent on the transformation machinery. In Figure 3 we provide a model that fits our observations (**Fig. 3**).

**SFig.4:**
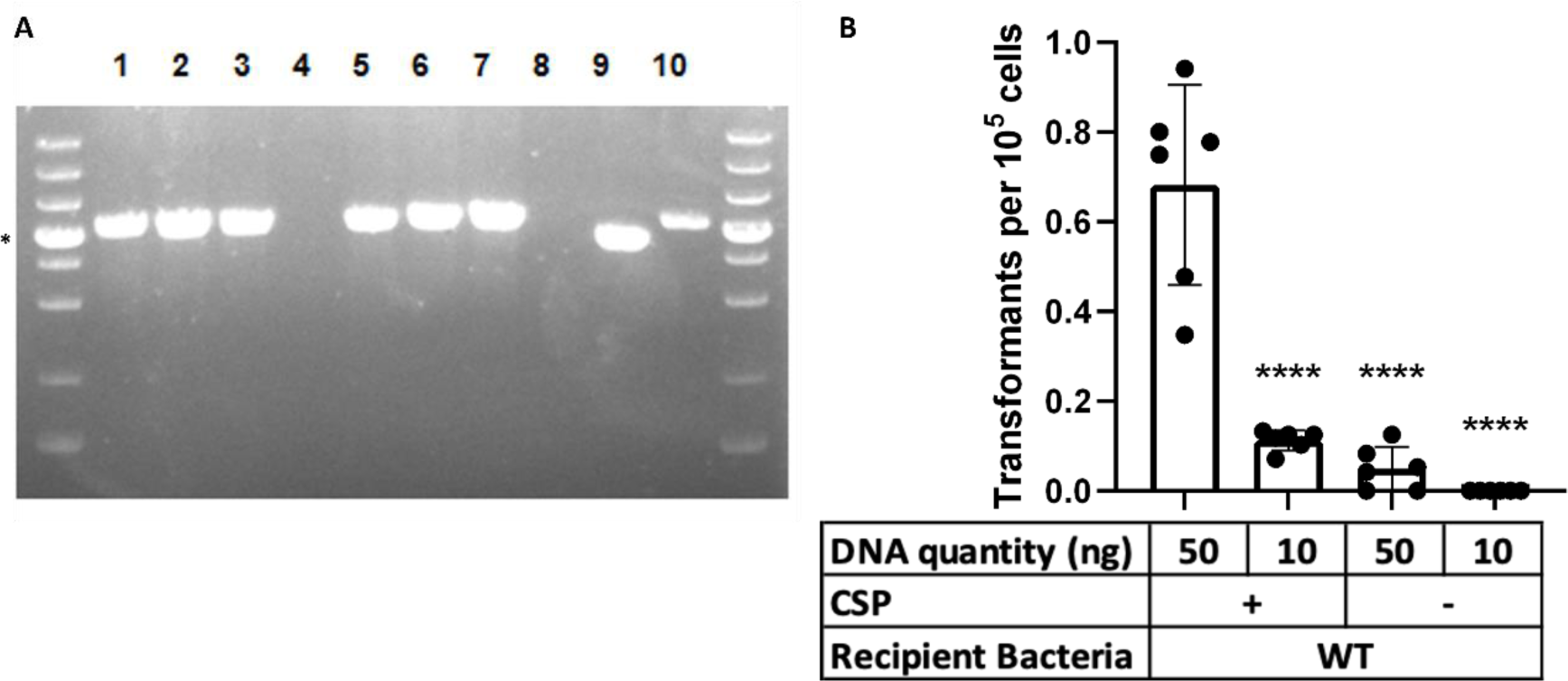
pEVs mediate horizontal gene transfer. **(A)** Three colonies from the transformation plates were grown in rich media overnight and used as a PCR template to check for presence of genes encoding spectinomycin resistance. Every colony produced amplicons of the appropriate size. The templates are as follows: Lanes 1-3 and 5-7: transformation colonies; lane 4: growth media; lane 8 no template; lane 9: gDNA from wild-type R6 SpecS; and lane 10: genomic data from donor bacteria R6-SpecR. Asterisk (*) indicates 5,000 base pair marker on the GeneRuler 1 kb Plus DNA Ladder (Invitrogen). **(B)** D39 cells (SpecS background) were exposed to pEV DNA from a D39-SpecR strain. Transformations were performed with and without CSP. Bars represent mean ± SEM with dots overlayed within a bar representing a data point from each independent experiment (n=6, **** adjusted p-value < 0.0001 for Dunnett’s multiple comparison test).

**Figure 3.**
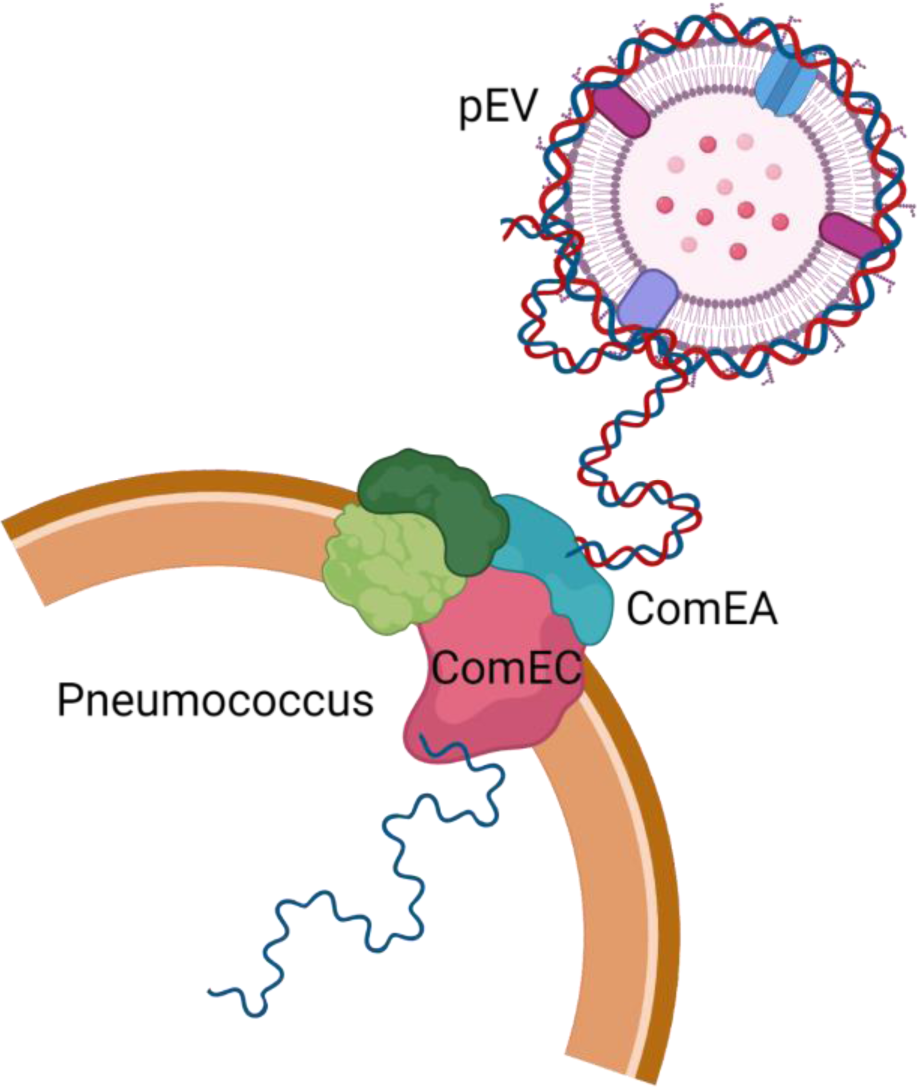
Working model of the transfer of pEV DNA to the transformation machinery of pneumococcal cells. DNA is associated with the extracellular surface of pEVs and can enter pneumococcal cells. Entry requires activation of competence by the competence stimulating peptide as well as a functional transformasome.

## Discussion

In this study, we demonstrate that pEVs are capable of mediating HGT. Our data support a model where pEVs deliver DNA to the cell via the transformation machinery and not through EV-fusion to the recipient cell membrane, nor by EVs transport into the recipient cell. This model raises multiple questions. First, what is the mechanism by which DNA associates with the outer surface of pEVs? We have established that the DNA matches that of producing cells, thus we speculate that pEVs either capture DNA that has been released from the cell or DNA is actively incorporated from the cytosol. In addition, there are questions regarding the binding dynamics of DNA to pEVs. What are the rates of association/dissociation from the pEV surface, and what are the molecular interactions tethering DNA to pEVs? These questions are all the subject of our ongoing investigations into pEV DNA.

An overarching question from Gram-positive EVs in general, and pEV specifically, is how they traverse the thick cell wall and why do we observe the range in their sizes. Brown et al, 2015 has put forward three models for release of pEVs (22). These models are not mutually exclusive, and all presume that pEVs bud from the plasma membrane. In the first model, pEVs are forced through pores in the cell wall by turgor pressure. In the second model, the cell wall is enzymatically modified to allow pEV release. In the third, channels present on the cell wall guide pEV release. Each of these models allow for variable pEV sizes dictated by pore or channel size, or the extent of enzymatic modification of the cell wall. Further, the measured range in pEV size may reflect not only the original size of these vesicles but also fusion and budding events post-pEV formation, which is consistent with our cryo-EM images.

Our findings do not exclude the possibility that pEVs may also deliver their contents directly to recipient cells. *Bacillus subtilis*, another Gram-positive bacterium, has been shown to receive protein content and integrate it into its cell envelope (36). Further, vesicles from the Gram-negative *A. baylyi* can deliver DNA to *E.coli* that are not competent, consistent with another means of EV-associated gene transfer (21). Thus, bacterial EVs may deliver material to bacteria in multiple ways.

While there is no prior evidence of EV-mediated HGT in a Gram-positive bacterium, in Gram-negative *A. baylyi* there is evidence that outer membrane vesicles can interact with the transformation machinery (21). Specifically, in this case, intra-species transfer of DNA was observed. However, experiments with transformation-deficient mutants of *A. baylyi* suggested that the competence locus was required for DNA transfer. These results suggest that the link between pEVs and transformation may be a widespread phenomenon, observed in both Gram-negative and Gram-positive bacteria.

## Material & Methods

### Bacterial strain selection and culture growth

Unless otherwise noted, the experiments were performed with *S. pneumoniae* R6 or D39 and mutants generated in these backgrounds. WGS experiments used pEVs purified from *S. pneumoniae* R6, ATCC700669-NP and B1599 (33). Bacteria were grown from frozen stocks by streaking on TSA agar plates containing 5% sheep blood, incubated overnight at 37°C in 5% CO_2_. Planktonic cultures were grown by picking colonies from TSA agar plates and inoculating them into Columbia broth at 37 °C in 5% CO_2_.

### Generation of mutant strains R6 and D39 Spec insertion and R6 *Δ*comEA/comEC

The R6 and D39 spectinomycin-resistance (SpecR) strains and R6 *ΔcomEA/comEC* strain in this work were constructed by inserting an antibiotic resistance cassette into the genome by homologous recombination. For the SpecR strains, a spectinomycin cassette was amplified from plasmid pr412 (a gift from Dr. Donald Morrison) and inserted in the intergenic region between *spr0515/spr0516* (*spd0511/spd0512*). The R6 *ΔcomEA/comEC* utilized a kanamycin resistance cassette from the Janus cassette inserted in the location of the *comEA* and *comEC* genes (*spr0856* and *spr0857)*. For all constructs, ∼2000 base pair length flanking segments from these specific regions were amplified using primers listed in **Table S2** and fused to the antibiotic resistance cassette by either restriction digestion and ligation (BamHI, XmaI, T4 DNA ligase) for the spectinomycin cassette insertion or by Gibson assembly for the kanamycin cassette insertion. These constructs were then transformed into *S. pneumoniae* strain R6 or D39. The recipient bacteria were grown in Columbia broth to an optical density of 0.05, then mixed with the construct and CSP1 ‘EMRLSKFFRDFILQRKK’ was added to 0.125 µg/mL final concentration. The mixture was incubated at 37°C for 2 h, then plated on Columbia agar plates with either spectinomycin (100 µg/mL) or kanamycin (150 µg/mL). The transformations were incubated at 37°C in 5% CO_2_ overnight. Colonies were picked and confirmed by PCR and Sanger sequencing.

### pEV Isolation

Pneumococcal cultures were grown in Columbia Broth to mid-to-late log phase (OD_600_ between 0.650 – 0.950), as previous studies demonstrated high yields at that stage of the growth phase (27). pEVs were isolated by size exclusion chromatography as described in previous work (28, 37). Conditioned media was centrifuged at 10,000-20,000 *x g* for 20 min at 4°C to pellet the bacterial cells. The supernatant was filter-sterilized by passing through a 0.22 µM Millipore filter (VWR 500 ml filtration units) and stored at 4°C. The supernatant was concentrated 500-fold using 100 kDA molecular weight cut-off filters (Thermo Scientific Pierce, Rockford, IL) that were centrifuged at 2,500 *x g* at 4°C until appropriately concentrated. The pEVs were isolated from the concentrated supernatant via mini-SEC using 1.5 x 12 cm mini-columns (Econo-Pac columns; bio-Rad, Hercules, CA) loaded with a 10 mL packed volume of Sepharose CL-2B (Sigma-Aldrich, St. Louis, MO), equilibrated with PBS. The concentrated supernatant was loaded onto the column, the eluent was PBS, and the samples were eluted in 1 mL fractions (pEVs elute in Fractions 4 and 5). Fraction 4 was collected, quantified, and either used immediately (within 24 hours) or stored at −80 °C for long-term storage.

### Nanoparticle Tracking Analysis (NTA)

The pEV particle size and count distribution was quantified by nanoparticle tracking analysis (NTA) using either a NanoSight 3000 (Malvern Panalytical, Malvern, UK) or a ZetaView (Particle Metrix). Each instrument automatically quantified the particle size, distribution, and concentration using the NTA software (version 3.4, Malvern Panalytical, Malvern, UK; ZetaView version 8.05.12, Particle Metrix GmbH, Ammersee, Germany).

### Protein quantification

The protein content of pEVs were quantified using Pierce BCA kit according to the manufacturer’s instructions (BCA Protein Assay Kit, Thermo Fisher).

### DNA quantification

DNA was quantified by spectrometry (NanoDrop 2000), qPCR, and PicoGreen dye. The qPCR reactions included: Sybrgreen with ROX, primers specific for the *S. pneumoniae* R6 16S rRNA gene, and R6 genomic DNA standards (range: 0.01-10 ng/µL) or pEV samples. The qPCR program (7300 system Applied Biosystems Sequencing Detection Software 1.4), primer sequences F: ACCCGAAGTCGGTGAGGTA R: CCAAATCATCTATCCCACCTT. The PicoGreen dye reactions included a 1:200 dilution of PicoGreen dye in PBS and either phage DNA standards (range: 0.002-2 ng/µL) or pEV samples according to the manufacturer’s instructions (Quant-iT PicoGreen dsDNA Kit; Invitrogen).

### PCR Amplification and Gel Electrophoresis

PCR amplifications were performed using the NEB Q5 high fidelity enzyme, pEVs as the experimental DNA template and gDNA as the control DNA template, and primers in Table S1. Amplicons were loaded onto a 1% agarose gel with a 1 kb Gene Plus Ruler ladder for gel electrophoresis.

### DNase Treatment and Lysis Assay

The pEVs underwent a series of treatments to determine the localization of DNA. At each treatment step, including the untreated pEVs, a sample was saved for PCR amplification and subsequent gel electrophoresis. First, pEVs were treated with 1 U Turbo DNase at 37°C for 30 minutes. Then, the DNase was inactivated by the addition of 5 µM EDTA (f.c.) for 10 minutes at 75 °C. Next, the pEVs were lysed with 1% triton (f.c.) at 65°C for 10 minutes. After the DNase inactivation step, an aliquot was treated with 50 ng of genomic DNA and incubated at 37°C for 30 minutes to ensure the DNase inactivation was complete. After the lysis treatment, an aliquot was treated with 50 ng of DNA and incubated at 65°C for 10 minutes to ensure the triton did not impede the subsequent PCR. The negative control was 1x PBS, which is the solution used for pEV elution from SEC. PCR amplification was performed using primers specific for the *S. pneumoniae* R6 *gapdh* gene (**Table S1**) and the amplicons underwent gel electrophoresis (1% agarose) for DNA migration and subsequent imaging.

### pEV staining by DiD and PicoGreen

1 µL of Vybrant DiD lipophilic dye (ThermoFisher) was added to 1 mL of pEVs and incubated at 37 ° C for 1 hour. PicoGreen (Invitrogen) was added in a 1:200 dilution and incubated at room temperature for 5 minutes. pEVs were concentrated in a Vivaspin 500 µL 100,000 MWCO concentrator (Sartorius) and washed with 5 mL of PBS to remove excess dye before resuspension in 250 µL of PBS.

### Single particle imaging, co-localization and co-diffusion analysis

Simultaneous pEV and DNA single particle imaging was performed via dual-camera spinning disc confocal microscopy. Images were acquired using a motorized Nikon Ti-2E microscope combined with a CREST X-Light V3 spinning disk unit outfitted with beam splitting optics and two cameras. The microscope was driven by Nikon Elements software and samples were imaged at 25° C in glass-bottomed 96-well plates (Mattek) using a 100x silicone immersion objective (Nikon Plan Apochromat, NA 1.35). The source of illumination was an LDI-7 Laser Diode Illuminator (89-North), and PicoGreen (Invitrogen) fluorescence and Vybrant DiD lipophilic dye (ThermoFisher) fluorescence were simultaneously excited at 488 nm and 635 nm respectively. A 565 nm dichroic mirror was used to direct the green channel to a back-thinned sCMOS camera (Hamamatsu Orca Fusion BT) and the far-red channel to a second sCMOS camera (Hamamatsu Orca Fusion). Single-plane images were acquired without delay using a 50 ms exposure time to enable particle tracking.

For co-localization analysis using Ripley’s cross-L function, images were first pre-processed by performing a rolling-ball background subtraction, followed by image filtering to remove noise (minimum, median, and Gaussian filters were applied to the pEV channel, while a Gaussian filter was applied to the DNA channel). Thresholds were then applied to each channel to generate binary masks of the pEV and DNA particles. The centers of the particles were defined using connected components analysis. The resulting 2D point pattern of DNA and pEV particle centers was analyzed using Ripley’s cross-L function, which quantifies the density of points at different spatial scales with respect to the (fixed) spatial distribution of a separate point pattern. We treated the DNA particle centers as the fixed-point pattern – thus, the heuristic interpretation of the cross-L function is the local density of pEV’s with respect to the distribution of DNA particles. We used a Poisson process for point generation as the null model for the pEV distribution, and a null acceptance region was derived from 1000 Monte Carlo simulations, as implemented in the spatstat package in R (38). Data points which lie above the acceptance region thus exhibit greater clustering of pEV’s to DNA particles than would be expected based on chance. We note that due to lateral chromatic aberration, DNA particles and pEV’s were not perfectly registered. Thus, for very small spatial scales (<0.1 µM), the observed cross-L function overlaps with the null acceptance region (**Fig. 1C-D**).

For co-diffusion analysis, non-stationary pairs of pEV’s and DNA particles were isolated and the centers of the particles were identified and tracked using the Crocker-Grier algorithm, as implemented by the trackpy package in Python (39, 40). Diffusion coefficients for pEV’s were computed based on a power-law fit to their squared-displacements/lag-time profiles in log-space. The Fréchet distances between the pairs of pEV and DNA particle trajectories were computed after registration. The Fréchet distance is heuristically defined as the minimum cord-length sufficient to join a point traveling forward along one curve with a point traveling forward along a second curve. Thus, it is a metric for the similarity between trajectories (large values correspond to greater divergence). Using the measured diffusion coefficients for each pEV particle, 10 Brownian trajectories were simulated starting from the same origin as the corresponding DNA particle (**Fig.S2**). Simulations were run for the same timespan as the corresponding data, and Fréchet distances were computed, resulting in the “Random” distribution displayed in **Fig. 1E**.

### Genomic and pEV DNA extraction

DNA was extracted from pneumococcal bacterial samples and from pEV samples. To prepare the bacterial samples, each strain was grown to mid-log growth phase, then centrifuged at 4,500 *x g* for 10 minutes to pellet the bacteria from the supernatant. The supernatant was discarded, and the cells were frozen at −20°C for future use. The cells were resuspended and thoroughly mixed in 460 µL of 1x TE buffer. For the pEV sample, an equal volume of SEC-purified pEVs were used. Each sample was then treated the same for DNA extraction.

Each sample was treated with the addition of 10 µL of lysozyme (100 µg/ml) and 10 µL of mutanolysin (200 µg/mL) and an incubation at 37°C for 30 minutes. Next, 6.25 µL of RNase A (4 mg/mL) was added to the sample and was subsequently incubated at 37 °C for 5 minutes. After five minutes, 10% SDS was added to the sample, which was then vortexed and incubated in a water bath for 37°C for 30 minutes. Then, 2.5 µL of 20 mg/mL Proteinase K was added to the sample, the sample was mixed thoroughly, and was incubated in a water bath at 55°C for 30 minutes. Then 90 µL of 5 M NaCl was added to the sample, the sample was mixed thoroughly, and 75 µL of pre-heated CTAB (0.275 M) /NaCl (0.07 M) was added to the sample, which was incubated at 65°C for 20 minutes. An equal volume of Phenol:Chloroform:Isoamyl Alcohol (25:24:1) was added to the sample, vortexed thoroughly, and centrifuged at 10,000 *x g* at room temperature for 10 minutes. The aqueous layer (approximately 600 µL) was transferred to a new tube and an equal volume of 100% isopropanol was added to the tube. The sample was gently inverted 50 times, then centrifuged at 10,000 *x g* at room temperature for 5 minutes. The supernatant was aspirated and 250 µL of cold 70% EtOH was added to the tube. The sample was vortexed to dislodge the pellet, then the sample was centrifuged at 10,000 *x g* at 4°C for 5 minutes. The supernatant was aspirated, and the pellet was air dried until the residual ethanol evaporated. Then 75 µL 1x TE buffer was added to the pellet and was incubated in a water bath at 55°C for 1 hour, quantified by spectrophotometry at 260/280, and then stored at 4°C until use.

### Whole genome sequencing

Extracted gDNA samples from the pEVs were sequenced in the Hartwell Center at St. Jude Children’s Research Hospital. Sequence libraries were prepared and barcoded using the Nextera kit and run on the Illumina Novaseq platform according to manufacturer guidelines. Trim Galore version 0.6.4 (https://www.bioinformatics.babraham.ac.uk/projects/trim_galore/) was used to remove adapters and low quality regions from paired-end reads. The clean reads were assembled with megahit (version 1.2.9) and QUAST (version 5.0.2) was used to compare each pEV assembly to its reference strain genome for coverage.

### pEV-mediated transformation

The transforming DNA quantities were either 10 ng or 50 ng per transformation reaction. The pEV samples were from either the R6-SpecR strain or the D39-SpecR strain. A genomic DNA control was performed for every pEV-mediated transformation, with identical quantities of DNA per transformation reaction. The recipient bacteria were either the R6 wild type strain, the R6 *ΔcomEA/comEC*, or the D39 wild type strain. The recipient bacteria were grown in Columbia broth to an optical density of 0.05. Transformation reactions occurred with the following variables: DNA source (pEV vs genomic DNA), DNA quantity per reaction (50 vs 10 ng), CSP1 +/-, recipient bacterial strain (WT vs *ΔcomEA/comEC*). The transformation cultures were incubated at 37°C for 1 h. After 1 h, two 100 µL quantities of the transformation culture were aliquoted for enumeration of total bacteria in the transformation culture and for quantifying transformation. The enumeration of total bacteria occurred by performing a 1:10 serial dilution using 1x PBS as the diluent. A 10 µL sample of the final dilution tube was plated on TSA agar with 5% sheep blood, grown overnight at 37°C in 5% CO_2_, and enumerated the following morning. The quantification of the transformation colonies occurred by spreading 100 µL of the transformation culture on Columbia agar and spectinomycin (100 µg/mL) growth media. The plates were incubated overnight. Colonies were picked, grown in Columbia broth at 37°C 5% CO_2_, and confirmed as transformants by PCR. Transformation Efficiency (TE) is calculated as transformed colonies/µg transformed DNA/DNA dilution = TE (cfu/µg DNA).

### Cryo-electron microscopy

Samples of pEVs were generally of low concentration when checked by negative-stain electron microscopy using a 2% solution of uranyl acetate. Consequently, cryo-EM was carried out with the automated imaging software “EPU” on a Titan Krios 3Gi equipped with a Selectris energy filter and a Falcon 4i direct electron detecting camera so that enough representative images could be collected of each sample. 3µL of each sample was was pipetted onto a freshly glow-discharged Quantifoil R2/1 grid (Quantifoil Micro Tools GmbH, Großlöbicha, Germany), then blotted to remove excess sample and plunge-frozen into a mixture of liquid ethane and propane cooled in a bath of liquid nitrogen using a Vitrobot Mk 4 cryo-plunger (41). Grids were then mounted in the Krios microscope and imaged at 300 kV accelerating voltage and a magnification of 81,000x corresponding to a pixel size of 1.5 Ångstroms at the sample. The energy slit width used was 10 eV, and the Falcon 4i camera was operated in electron counting and movie modes with a total recorded dose of approximately 30 e/Å^2^. Drift was corrected from movies with MotCorr2 software and images were examined by for the presence of pEVs (42). All equipment and the EPU software were supplied by Thermo Fisher Scientific (Thermo Fisher Scientific, Waltham, Massachusetts, USA).

## Acknowledgements

This work was generously supported by a grant from the Shurl and Kay Curci Foundation. Further, graduate student support was generously provided by the Shurl and Kay Curci Foundation for SSY, the NSF Graduate Research Fellowship to SWL, and the NIGMS Predoctoral Institutional Research Training Grant T32 under award 5T32GM133353-04 to SC and BES. The University of Pittsburgh Titan Krios microscope were supported by the Office of the Director, National Institutes of Health, under award S10 OD025009. The work also received support from the Department of Biological Sciences at Carnegie Mellon University. The content is solely the responsibility of the authors and does not necessarily represent the official views of the National Institutes of Health. The funders had no role in study design, data collection and analyses, decision to publish, or preparation of the manuscript.

---

**Table S1.**
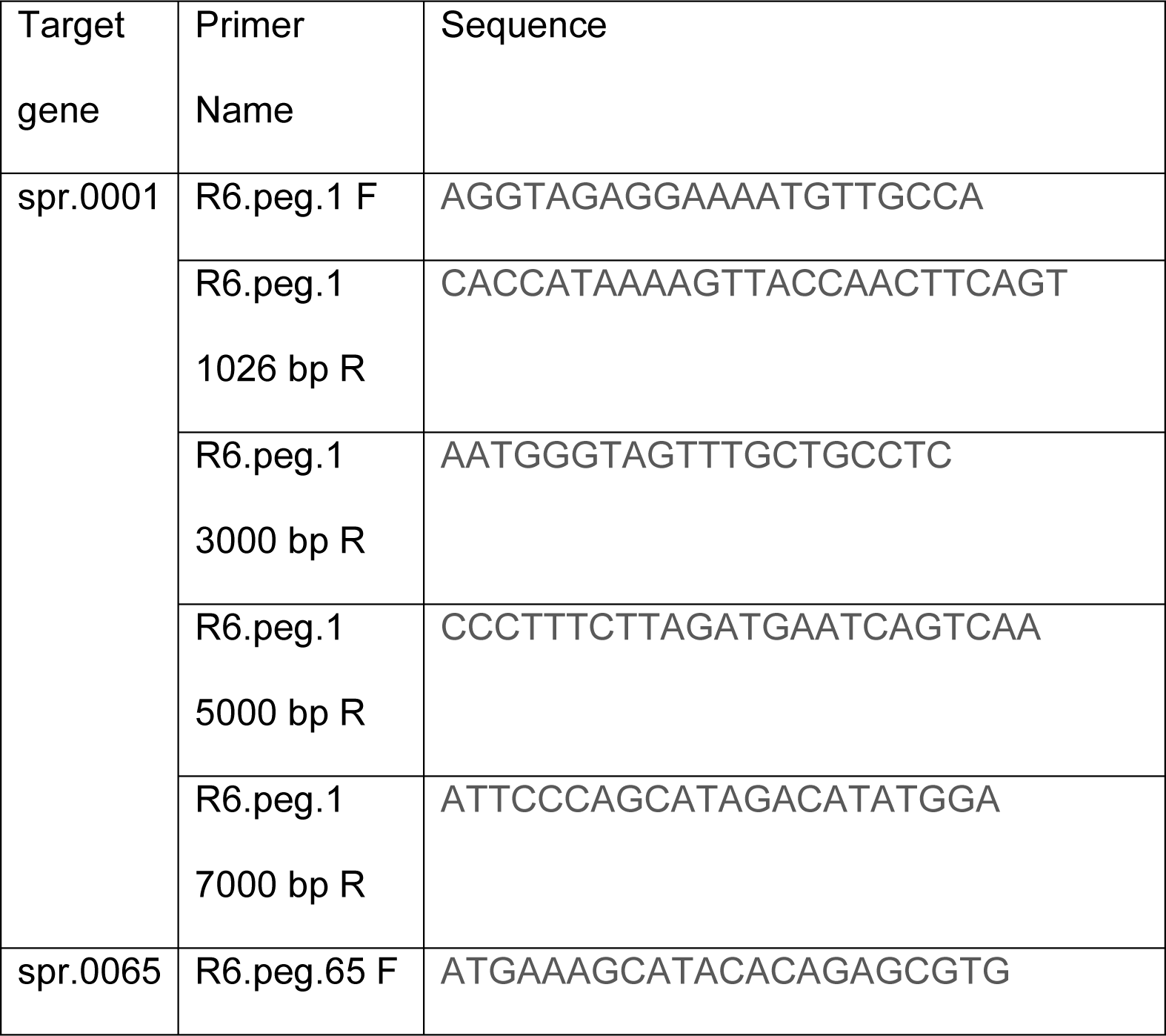
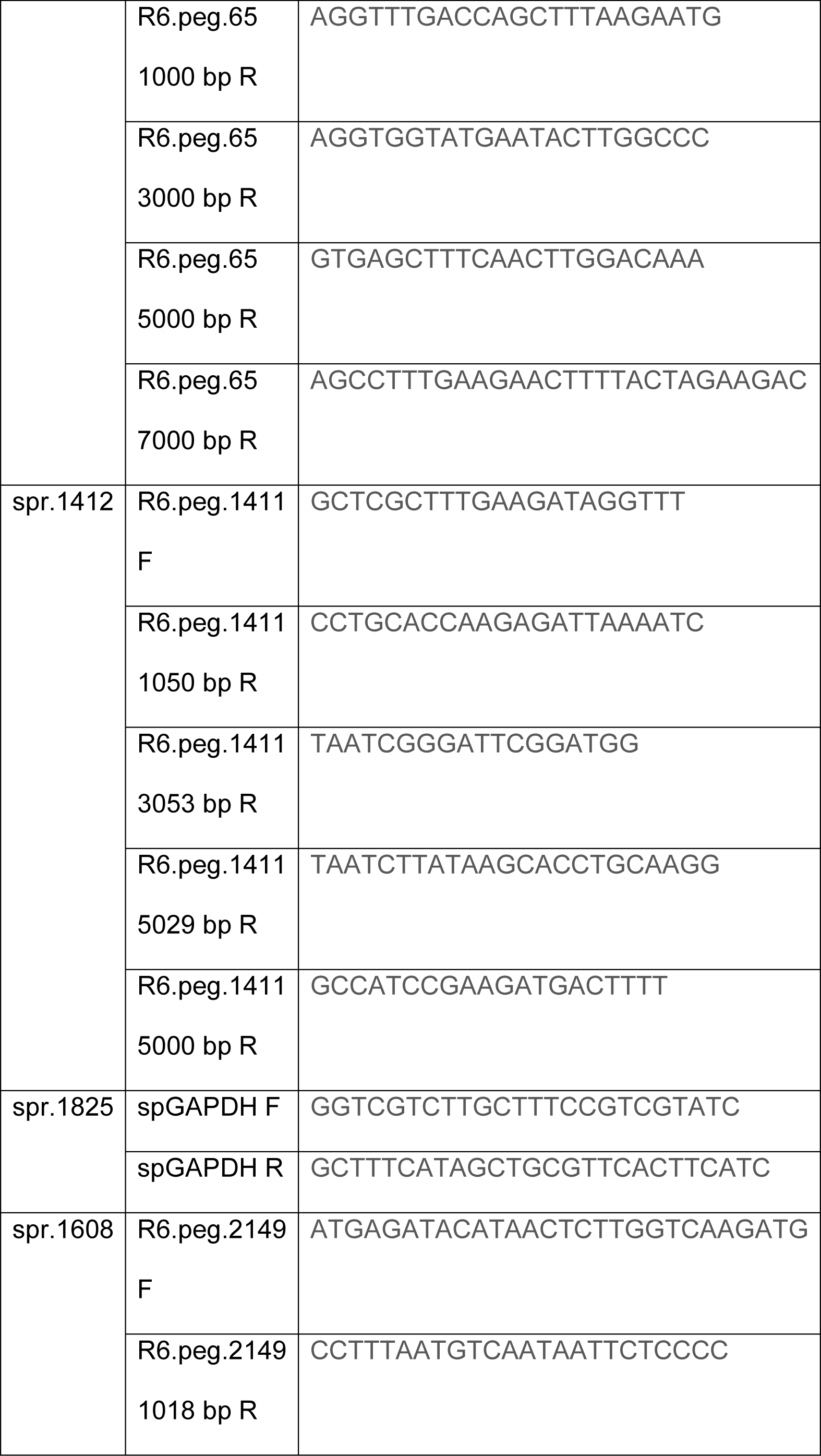
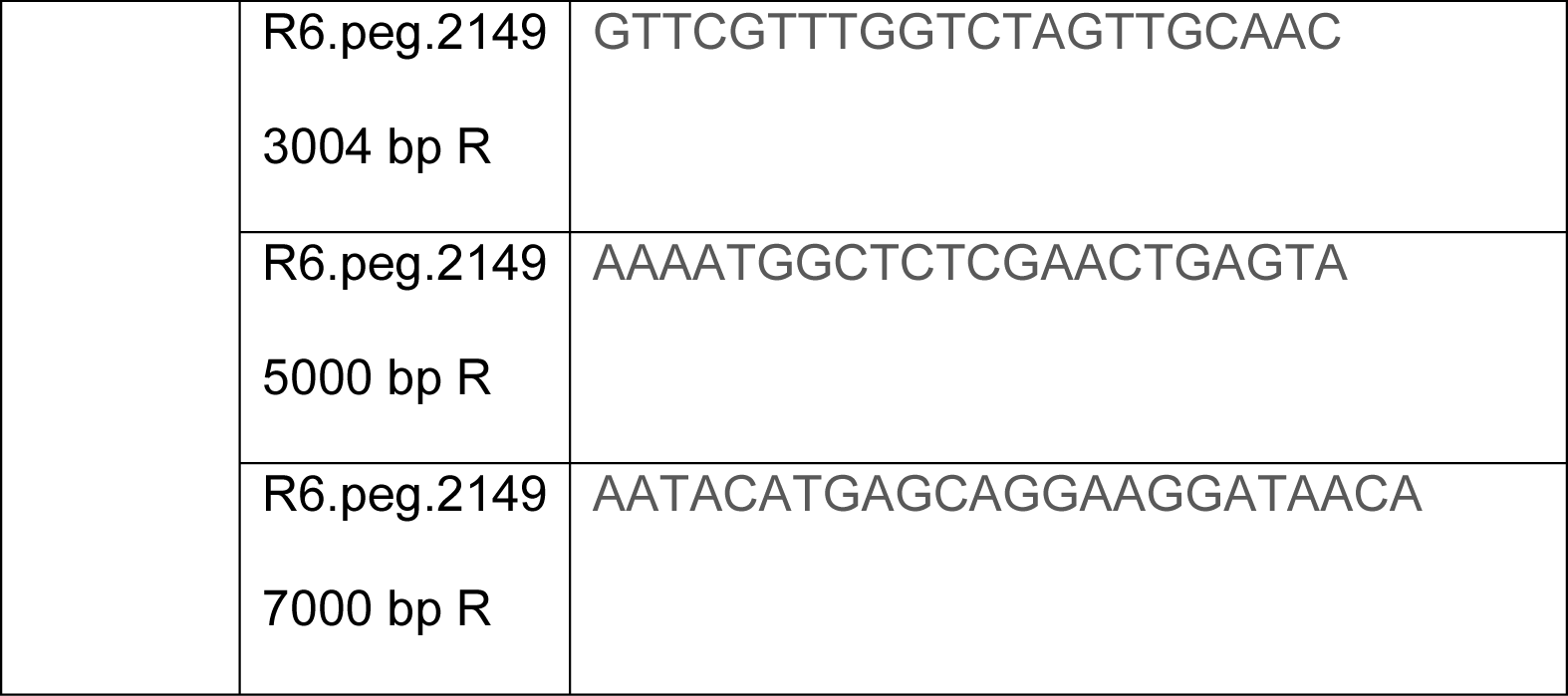
Primers for pneumococcal regions.

**Table S2.**
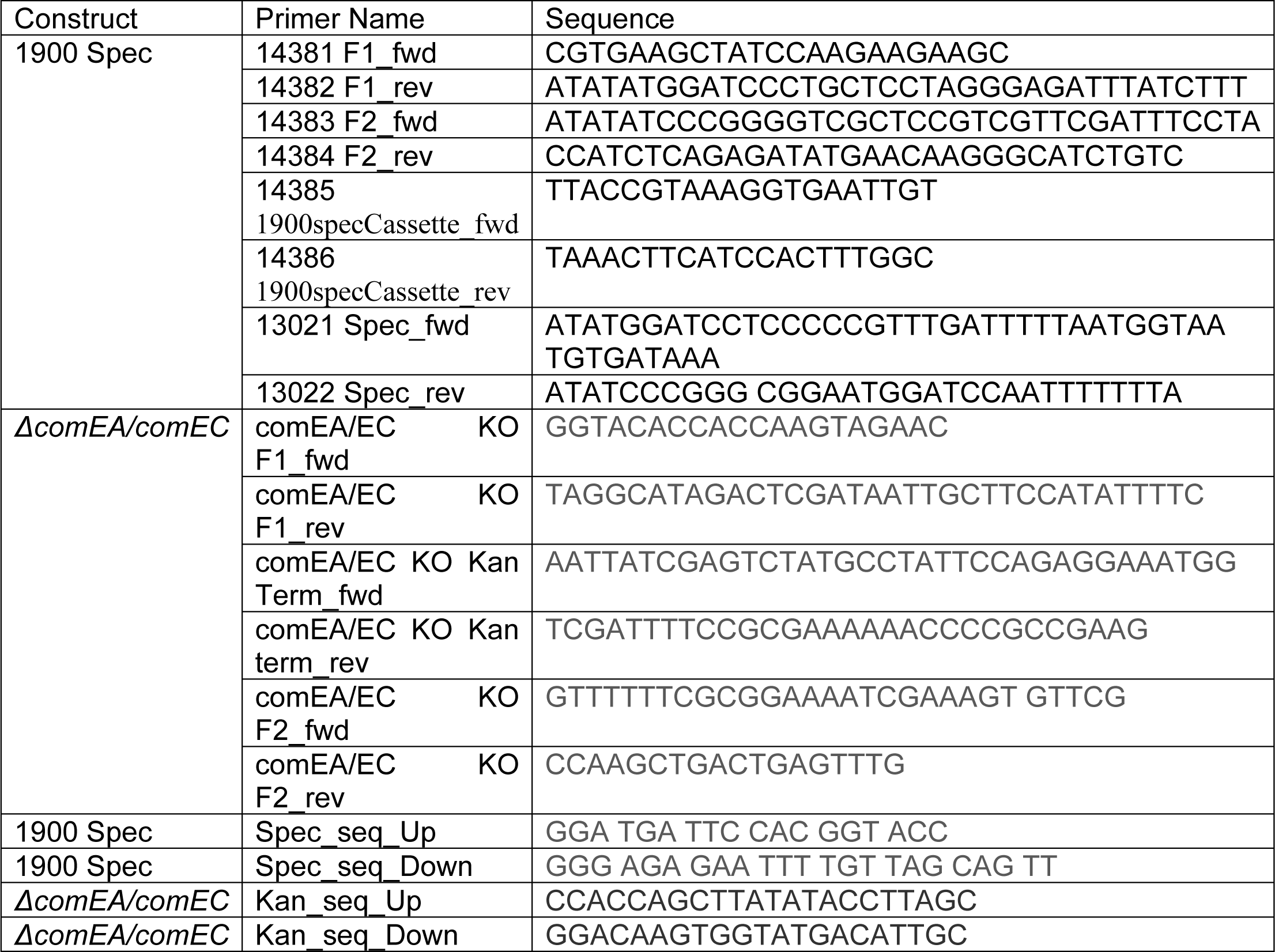
Gibson assembly primers and sequence check primers.

## Notes

### Competing Interest Statement

The authors have declared no competing interest.

